# Employing Artificial Neural Networks for Optimal Storage and Facile Sharing of Molecular Dynamics Simulation Trajectories

**DOI:** 10.1101/2024.09.15.613125

**Authors:** Abdul Wasim, Lars V. Schäfer, Jagannath Mondal

**Affiliations:** Tata Institute of Fundamental Research Hyderabad, Telangana 500046, India; Center for Theoretical Chemistry, Faculty of Chemistry and Biochemistry, Ruhr University Bochum, 44780 Bochum, Germany

## Abstract

With the remarkable stride in computing power and advances in Molecular Dynamics (MD) simulation programs, the crucial challenge of storing and sharing large biomolecular simulation datasets has emerged. By leveraging AutoEncoders, a type of artificial neural network, we developed a method to compress MD trajectories into significantly smaller latent spaces. Our method can save up to 98% in disk space compared to xtc, a highly compressed trajectory format from the widely used MD program package GROMACS, thus facilitating storage and sharing of simulation trajectories. Atom coordinates are very accurately reconstructed from compressed data. The method was tested across a diverse sets of biomolecular systems, including folded proteins, intrinsically disordered proteins, phospholipid bilayers, proteinligand complexes, large protein complexes and membrane-bound protein systems. The reconstructed trajectories demonstrated consistent accuracy in recovering key biophysically relevant properties for proteins, lipids and composite systems. The compression efficiency was particularly beneficial for larger systems. This approach enables the scientific community to efficiently store and share large-scale biomolecular simulation data, potentially enhancing collaborative research efforts. The workflow, termed “compresstraj”, is implemented in PyTorch and is publicly available at https://github.com/SerpentByte/compresstraj, offering a practical solution for handling the increasing volumes of data generated in biomolecular simulation studies.

## Introduction

Molecular Dynamics (MD) simulations are increasingly used to explore and understand complex biomolecular systems.^1^ MD trajectories, particularly for large simulation systems such as biomolecules, can generate impressive amounts of data ranging from hundreds of gigabytes to tens of terabytes. The storage and sharing of such data is understandably challenging. Since researchers involved in MD simulations have different goals, perspectives, and understandings of the systems they study, these trajectories are often underutilized regarding the information that could be extracted from their analysis. An effective distribution of these data sets would enable multifaceted exploration, increasing the efficiency of knowledge mining. The biomolecular simulation community has greatly benefited from established databases such as PDB and SwissProt. However, a significant gap remains: the lack of a unified database for storing and sharing simulation trajectories. Efforts such as MDDB,^2–4^ mdCATH^5^ and NMRlipids^6^ aim to fill this void, but the size of the data presents a substantial challenge. Platforms like Zenodo, OSF, and Figshare host hundreds to a thousand terabytes of data,^7^ but managing and accessing them is time- and resource-intensive. Furthermore, due to challenges faced in uploading such huge amounts of data, only a fraction of the existing simulation data is actually uploaded. Hence, a unified database could revolutionize data accessibility.

To address these challenges, existing methods exploit temporal coherence, molecular structure, or functional approximations to compress molecular dynamics (MD) trajectories.^8–14^ PCA-based approaches, such as essential dynamics compression,^8^ reduce dimensionality by projecting trajectories onto dominant collective modes, retaining major motions while discarding high-frequency fluctuations. Predictive coding strategies leverage frame-to-frame continuity using linear predictors,^10^ piecewise-linear segmentation as in HRTC,^11^ or bonded-rotation models that improve fidelity by accounting for local structural constraints.^12^ Coarse-grained methods, such as the one reported by Cheng et al.,^9^ reduce data by eliminating degrees of freedom, enabling fast compression and decompression. However, their compression capacity is inherently limited to approximately one-third of the original data size (∼67% compression) and, importantly, the compression is not lossless. More recently, adaptive frameworks like MDZ^13^ combined delta coding, quantization, and entropy optimization with configurable error bounds and high throughput, achieving some of the best error-controlled compression ratios reported to date. Parallel efforts using NetCDF4/HDF5 with lossy quantization filters have demonstrated high compression with negligible energetic distortion, highlighting the practical relevance of precision-aware formats.^14^ In this study, we use AutoEncoders (AE)^15^ to encode each frame of a molecular dynamics trajectory into a compact latent space representation, which we treat as its compressed form.

AEs are self-supervised neural networks^15^ that learn to compress input data into a low-dimensional latent space and then reconstruct the original data from this representation. In an ideal scenario, a perfectly trained AE would reproduce outputs identical to its inputs. In practice, however, perfect reconstruction is unattainable, so the goal is to minimize the reconstruction error—the difference between the input and the output. Because the targets are identical to the inputs, the training process is termed self-supervised. The network autonomously learns to encode the input into a latent representation and decode it back to the original space.

AEs have been used in biomolecular simulations primarily to project high-dimensional data, such as distance matrices, onto lower-dimensional latent spaces.^16–18^ However, using coordinates as input to AutoEncoders is less common ^19^ due to their lack of special Euclidean group in 3 dimensions (SE(3)) invariance, a rotation or translation of the entire system results in a different set of coordinates although the system’s fundamental features remain unchanged. In this study, we directly use atomic coordinates to train AutoEncoders, employing the latent space as the compressed trajectory. We demonstrate that the latent spaces represent highly compressed MD trajectories, with particle coordinates being very accurately reconstructed upon decompression. This enables the generation of highly compressed MD trajectories that can be easily stored, shared, and quickly decompressed when needed. Importantly, the ML-reconstructed trajectories accurately recapitulate key relevant structural properties, in close agreement with the original trajectories. We also provide user-friendly Python scripts and libraries to enable straightforward application of the protocol.

## Methods

### Details on the systems used

We have utilized the following biomolecular systems for this study (Figure 1 and Table 1): chignolin,^20^ trpcage,^21^ and Hen Egg White Lysozyme (HEWL)^20^ as representatives of folded proteins of different sizes; Amyloid *β*40 (A*β*40)^22^ and ACtivator for Thyroid hormone and Retinoid receptors (ACTR)^20^ as representatives of intrinsically disordered proteins (IDPs); a DOPC bilayer as representative of lipid membranes (data reused from NMRlipids database entry 330);^6^ the L99A mutant of T4 Lysozyme with benzene as the ligand (T4L L99A + benzene)^23^ and cytochrome P450 with camphor (Cyt-P450 + camphor)^24^ as representatives of protein-ligand systems; the N-glycosylated SARS-CoV2 spike protein^25^ as a representative for a very large protein complex with amino acid modifications; CymA embedded in a PHPC bilayer^26^ as a representative of a membrane-bound protein system; and TIP3P water as a representative of water used in simulations.

**Figure 1:**
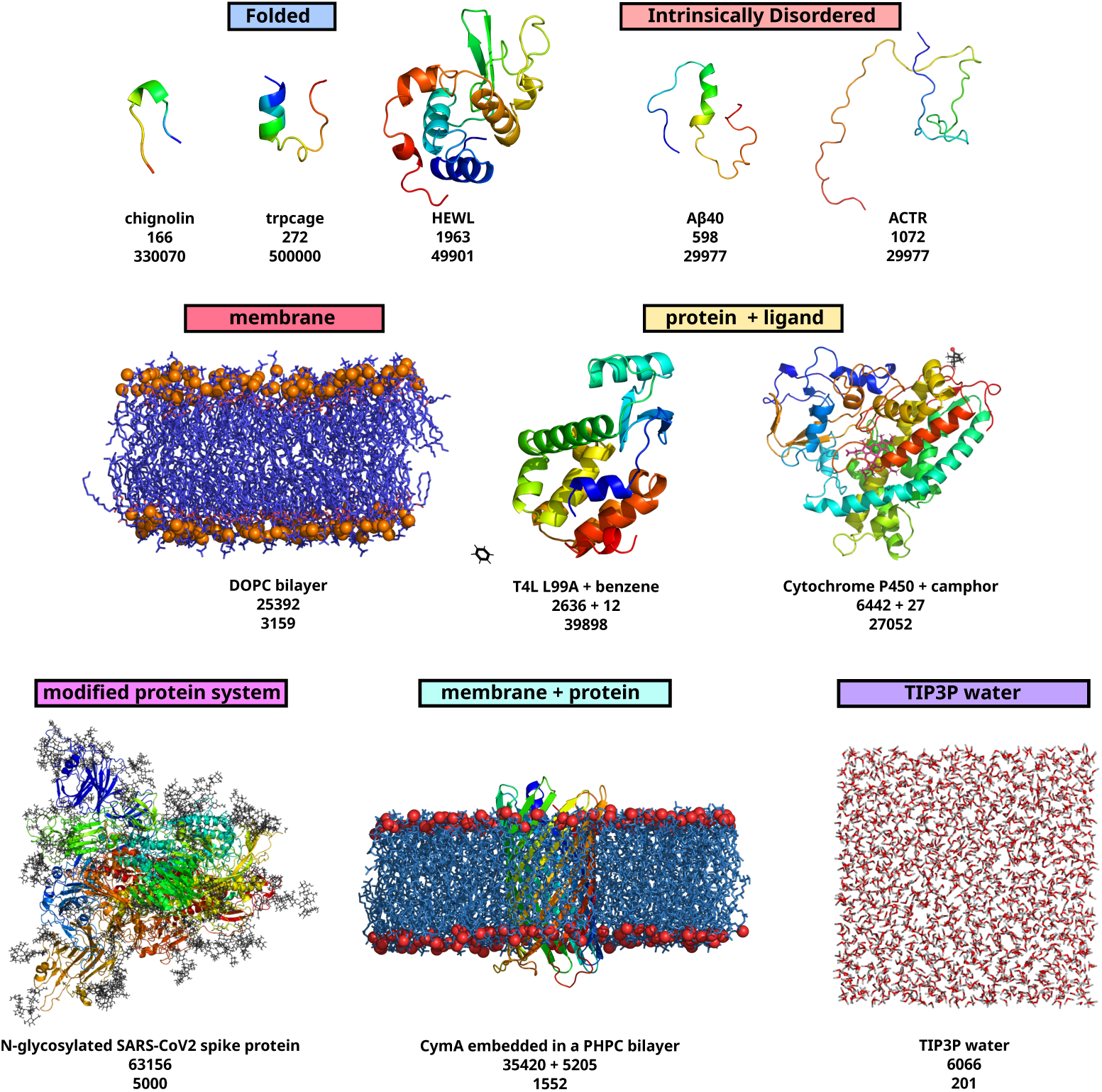
Representative conformations of the systems used in this study, along with the number of atoms (middle) and the respective number of frames in each trajectory (bottom). The data for folded and intrinsically disordered proteins were obtained from D. E. Shaw Research. ^20^ The DOPC bilayer was simulated in-house, with lipid heads shown in orange and tails in blue. The data for T4L L99A + benzene were obtained from, ^27^ with benzene depicted in black. The data for cytochrome P450 + camphor were obtained from, ^24^ with camphor shown in black and its oxygen represented as a red sphere. The data for the SARS-CoV-2 spike protein were obtained from, ^25^ with glycans depicted in black. The data for the PHPC membrane + CymA were obtained from, ^26^ with lipid heads shown in red and tails in blue. The data for TIP3P water were generated in-house, with water oxygen atoms shown in red and hydrogen atoms in grey.

**Table 1:** Details of different simulation systems and MD trajectories used in this work.

| **system** | **# particles** | **# frames** | **XTC size (MB)** | **reference** |
| --- | --- | --- | --- | --- |
| chignolin | 166 | 330070 | 226 | Ref.<sup>20</sup> |
| trpcage | 272 | 50000 | 55 | Ref.<sup>20</sup> |
| HEWL | 1963 | 49901 | 369 | Ref.<sup>20</sup> |
| Aβ40 | 598 | 29977 | 70 | Ref.<sup>20</sup> |
| ACTR | 1072 | 29977 | 123 | Ref.<sup>20</sup> |
| DOPC bilayer | 13824 | 3001 | 156 | NMRLipids<sup>6</sup> entry 330 |
| T4L L99A + benzene | 2636 + 12 | 39898 | 385 | Ref.<sup>23</sup> |
| Cyt-P450 + camphor | 6442 + 27 | 27052 | 636 | Ref.<sup>24</sup> |
| SARS-CoV2 spike | 63156 | 5000 | 1228 | Ref.<sup>25</sup> |
| PHPC bilayer + CymA | 35420 + 5205 | 1552 | 230 | Ref.<sup>26</sup> |
| water | 6056 | 7050 | 4.1 | this work |

These systems cover a broad spectrum concerning size and flexibility, ranging from flexible small peptides (chignolin) to folded miniproteins/globular proteins (trpcage, HEWL) via intrinsically disordered proteins (A*β*40, ACTR) to membranes (DOPC bilayer), protein interacting with ligand (T4L L99A + benzene, Cyt-P450 + camphor), modified amino acids and large protein complexes (SARS-CoV2 spike protein), membrane-protein systems (CymA in PHPC bilayer) and solvents (TIP3P water). The availability of long time-scale simulation trajectories of these systems (either kindly shared by D. E. Shaw Research or obtained from online databases or generated in-house) prompted their choice in the present investigation.

### Processing of the trajectories

The first step of the protocol is to choose one trajectory frame as a reference configuration. The particular choice of frame is arbitrary and doesn’t affect the results (see below). The trajectory is first aligned with the reference frame, and for each frame of the aligned trajectory the center of mass of the system is set to the origin. These transformations render the resulting coordinates pseudo SE(3) invariant, so that they can be used as input to a standard AE.

For a system with *N* particles, the trajectory has 3*N* Cartesian coordinates. For training the AE, we first scale the coordinates along each dimension using a mix-max scaler, with the minimum value set to 0 and the maximum value set to 1. The (*N,* 3) coordinates, for each frame, are then flattened resulting in a vector with 3*N* dimensions. All coordinates are then pooled. Thus, for *f* frames, the resultant dataset is (*f,* 3*N*). The scaled trajectory is then used as input to the AutoEncoder.

### Architecture of the AutoEncoder

The architecture of the AutoEncoder used in this work is consistent across all systems studied (Figure 2). Here, we utilize a dense AutoEncoder, meaning that only fully connected feed-forward layers are employed. There are two dense layers in both the encoder and the decoder. The number of neurons in each layer and the sequence of occurrences of layers can be summarized as (input, 4096, 1024, L, 1024, 4096, output), where L is the number of latent dimensions. The inputs are original, scaled coordinates and the outputs are reconstructed, scaled coordinates.

**Figure 2:**
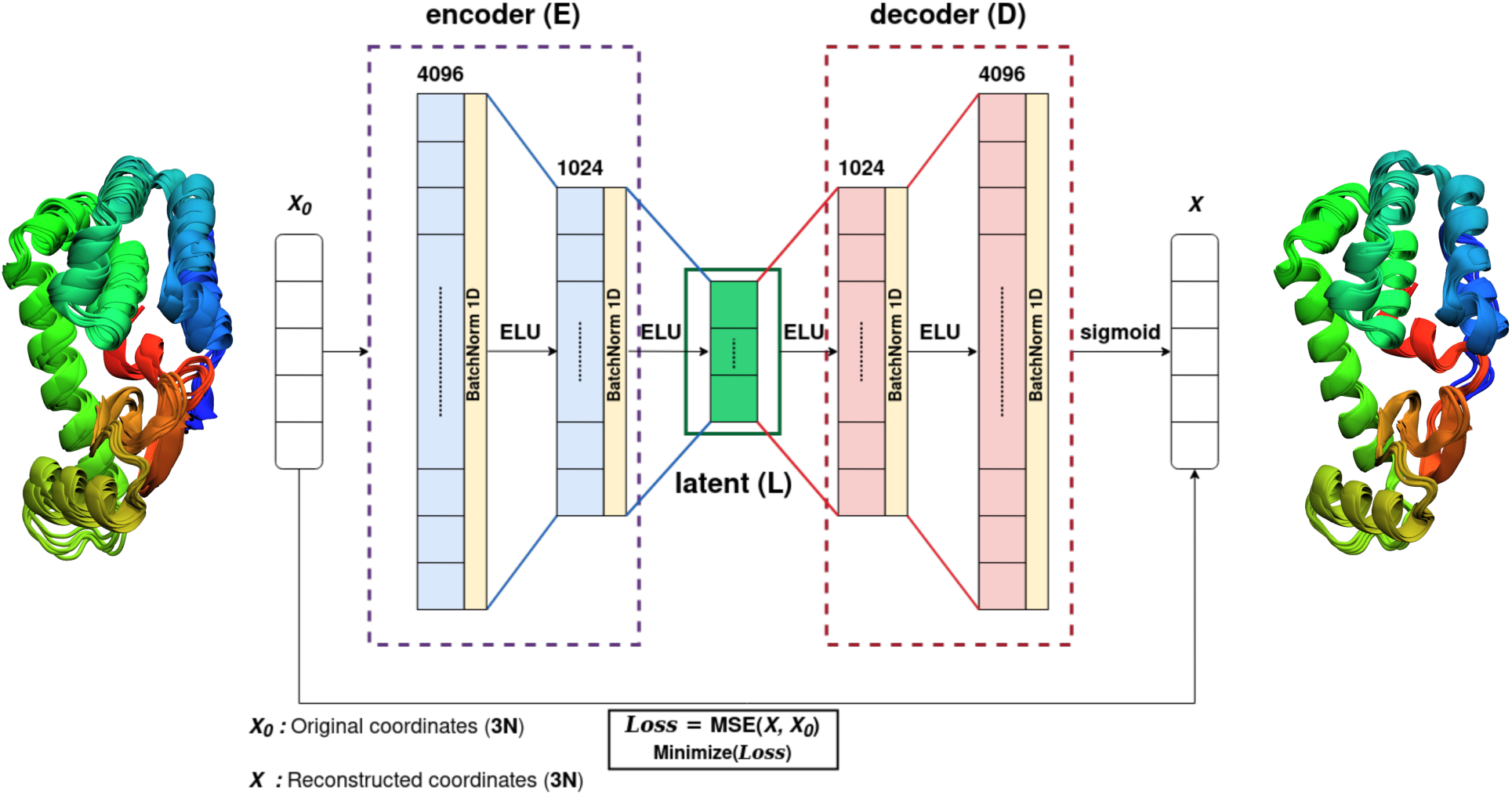
The architecture of the AutoEncoder used in this study. The input and the reconstructed input are *x*_0_ and *x*, respectively, and *L* is the latent space dimension. BatchNorm 1D is one dimensional batch normalization.

The models were built using PyTorch (see Figure 2 and *Supplementary Methods* for details on the architecture) and trained using PyTorch lightning. We use the exponential linear unit (ELU)^28^ activation function for all layers, except for the output layer, where a sigmoid activation function is applied. The models were trained using a root mean squared error (RMSE) loss function and the AdamW^29^ optimizer with the default hyper-parameters (*βs*=(0.9, 0.999), *ϵ*=1e-08, *λ*=0.01). Each model is trained for at least 2000 epochs (recommended for actual use cases). We fix all seeds for each training (we have used a seed value of 42 for training all models).

### Implementation of the algorithm and usage

To facilitate the preprocessing of trajectories and the training of AutoEncoders, we have developed a Python library named “compresstraj”, accessible via GitHub (https://github.com/SerpentByte/compresstraj). The repository includes ready-to-use Python scripts: one for compression, another for decompression, and one tailored for recombining fragmented systems. These tools are designed for out-of-the-box functionality upon library installation, streamlining the compression and decompression processes (see the GitHub repository for examples and further usage instructions).

#### Compression

For trajectory compression, the center of geometry of the unwrapped and aligned coordinates are first set to zero, followed by coordinate scaling between 0 and 1. The scaled coordinates are used as inputs to the AutoEncoders, as described in the *Methods* section. Once the system has been aligned with the reference conformation for a selected set of particles, the center of geometry is recorded for each frame. The coordinates are scaled between 0 and 1, flattened, and used to train an AutoEncoder. The default is to set the latent space of the AutoEncoder to 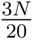 (or *c* = 20), where *N* represents the number of atoms/particles in the selected system or fragment (please see [MDAnalysis atom selection](https://docs.mdanalysis.org/stable/documentation_pages/selections.html) for selection commands). If 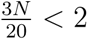, the code will automatically set the latent space dimension to 2, ensuring a minimum latent space size for the AutoEncoder.

Once the AutoEncoder model has been trained, the coordinates are projected onto the latent space using the encoder part of the AutoEncoder. This results in a compressed trajectory.

#### Decompression

The latent representations are first decoded into scaled coordinates using the decoder part of the AutoEncoder. These coordinates are then inverse-transformed to recover the original scale and, if a file containing the centers of geometry (COGs) is provided, translated accordingly. The resulting atomic positions are written to a molecular dynamics trajectory file, with the default output format being GROMACS’s compressed xtc format.^30,31^ This output format is configurable and can be changed to any format supported by the MDAnalysis library.^32^

Since the algorithm operates independently of whether the input is a complete system or a fragment, any system can be split into fragments and each fragment can be treated as an independent unit. Compression and decompression are performed individually on each fragment, which are then recombined using recompose.py to reconstruct the full trajectory. For example, systems such as T4L-L99A with benzene, Cytochrome P450 with camphor, and membrane-bound proteins such as cymA in a PHPC bilayer were processed by training separate models for each component and subsequently merging the outputs.

#### Resource Usage

All models were trained on an NVIDIA RTX 4090 Ti, except for the cymA embedded in PHPC bilayers and the glycosylated SARS-CoV-2 homo-trimer, which were trained on an NVIDIA L40S. The models are implemented in PyTorch with explicit CPU fallback, enabling training even in the absence of a GPU. While GPU acceleration significantly speeds up training—as is typical for deep neural networks—it is not strictly required. Importantly, compression and decompression are much faster than training and do not require a GPU. However, since training and compression are bundled together in the workflow, it may appear that compression requires a GPU, which is not the case.

The GPU memory usage for each tested system is summarized in Table 2, with all measurements recorded using a batch size of 256.

**Table 2:** GPU memory usage for training across systems of varying size.

| **System** | **Particles** | **GPU RAM usage (MB)** |
| --- | --- | --- |
| chignolin | 166 | 450 |
| trpcage | 272 | 516 |
| Aβ40 | 598 | 670 |
| ACTR | 1072 | 894 |
| HEWL | 1963 | 1320 |
| T4L L99A + benzene | 2648 | 1640 |
| TIP3P water | 6066 | 3314 |
| Cyt. P450 + camphor | 6469 | 4237 |
| DOPC bilayer | 13824 | 7875 |
| cymA | 40625 | 20252 |
| SARS-CoV2 | 63156 | 31252 |

Table 3 presents the GPU memory usage for the HEWL system under varying trajectory lengths and batch sizes. From these resource usage statistics, we conclude that the memory consumption does not increase with longer trajectories. However, larger batch sizes for the same system result in higher GPU memory usage.

**Table 3:**
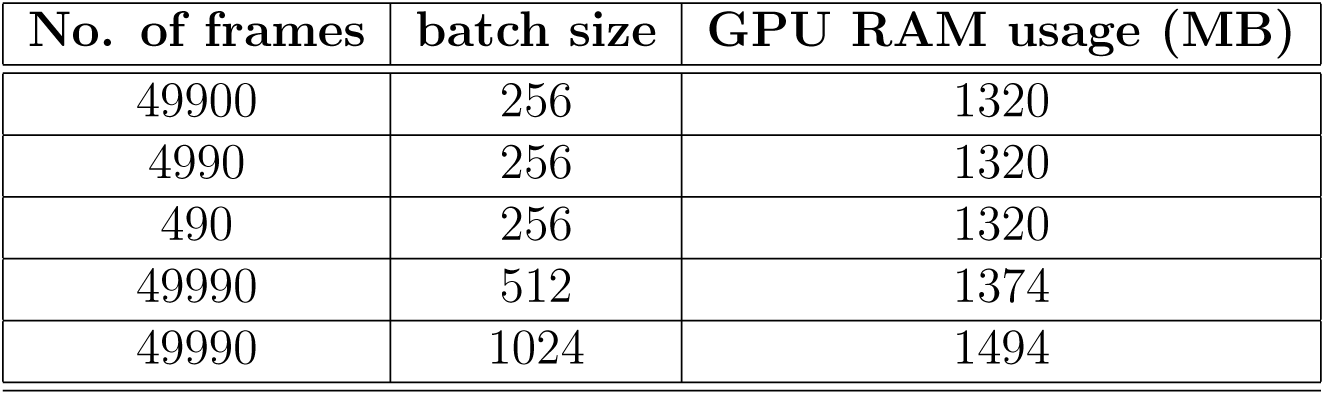
GPU memory usage for HEWL with varying trajectory length and batch size.

| **No. of frames** | **batch size** | **GPU RAM usage (MB)** |
| --- | --- | --- |
| 49900 | 256 | 1320 |
| 4990 | 256 | 1320 |
| 490 | 256 | 1320 |
| 49990 | 512 | 1374 |
| 49990 | 1024 | 1494 |

### Order parameters analysis of reconstructed trajectories

The quality of the decompressed trajectories reconstructed using the ‘compresstraj’ algorithm was assessed by comparing various biophysically relevant structural order parameters against the ground-truth (original) trajectory. In addition to basic comparisons, such as the root mean squared deviation (RMSD) of conformations and structural overlays, protein-specific properties – including Ramachandran plots, radius of gyration (*R_g_*), and C*_α_* contacts – were analyzed. Furthermore, lipid-related properties, including the density profiles of phospholipid head groups and lipid tail order parameters were characterized. For protein-ligand complexes, ligand binding profiles and their correlation with protein conformational changes were compared between the reconstructed and actual trajectories. All relevant details are provided in the respective subsections of the *Results and Discussion* section.

### List of Software

We have used only open-source software for this study. Snapshots were generated using PyMOL 2.5.4^33–35^ and VMD.^36^ Analyses were performed using Python^37^ and MDAnalysis.^32,38^ Figures were prepared using Matplotlib,^39^ Jupyter^40^ and Inkscape.^41^ All deep neural network models were built using Scikit-learn^42,43^ and PyTorch.^44^

## Results and Discussion

### Compressing Hen Egg White Lysozyme trajectory and effect of its reconstruction on Conformational ensemble

To gain a realistic idea of the extent of compression and the quality of reconstruction, we started our investigation of our AutoEncoder-based workflow on Hen Egg White Lysozyme (HEWL) (PDB ID: 1AKI), a folded protein with 129 residues that has served as a reference system for MD tutorials^45^ and benchmark studies.^20^

We trained the AutoEncoder on a trajectory with 49901 frames, which contained the three-dimensional Cartesian coordinates of all HEWL atoms, including hydrogens. Using the AE architecture detailed in the *Methods* section (see Figure 2) with atomic coordinates as input features, we trained the AE for 1000 epochs with a latent space dimension that is 20 times smaller than the input dimension (*c* = 20), using root mean squared error (RMSE) between scaled input and output as the loss function. Figure 3a) shows that the training loss drops rapidly, after which it changes very slowly. The distribution of RMSDs between original and reconstructed conformations is very narrow (Figure 3b) with a mean of 26 pm and a standard deviation of 1.9 pm. This shows that the reconstruction of the atomic coordinates from the compressed trajectory is very accurate. Furthermore, to ascertain whether the reconstructed images retain the secondary structure sampled in the original trajectory, we compare the Ramachandran plots obtained from the original (Figure 3c) and the reconstructed (Figure 3d) trajectories. As expected from the small RMSD values, these analyses show that the secondary structure is preserved upon reconstruction of the trajectories from the compressed latent space.

**Figure 3:**
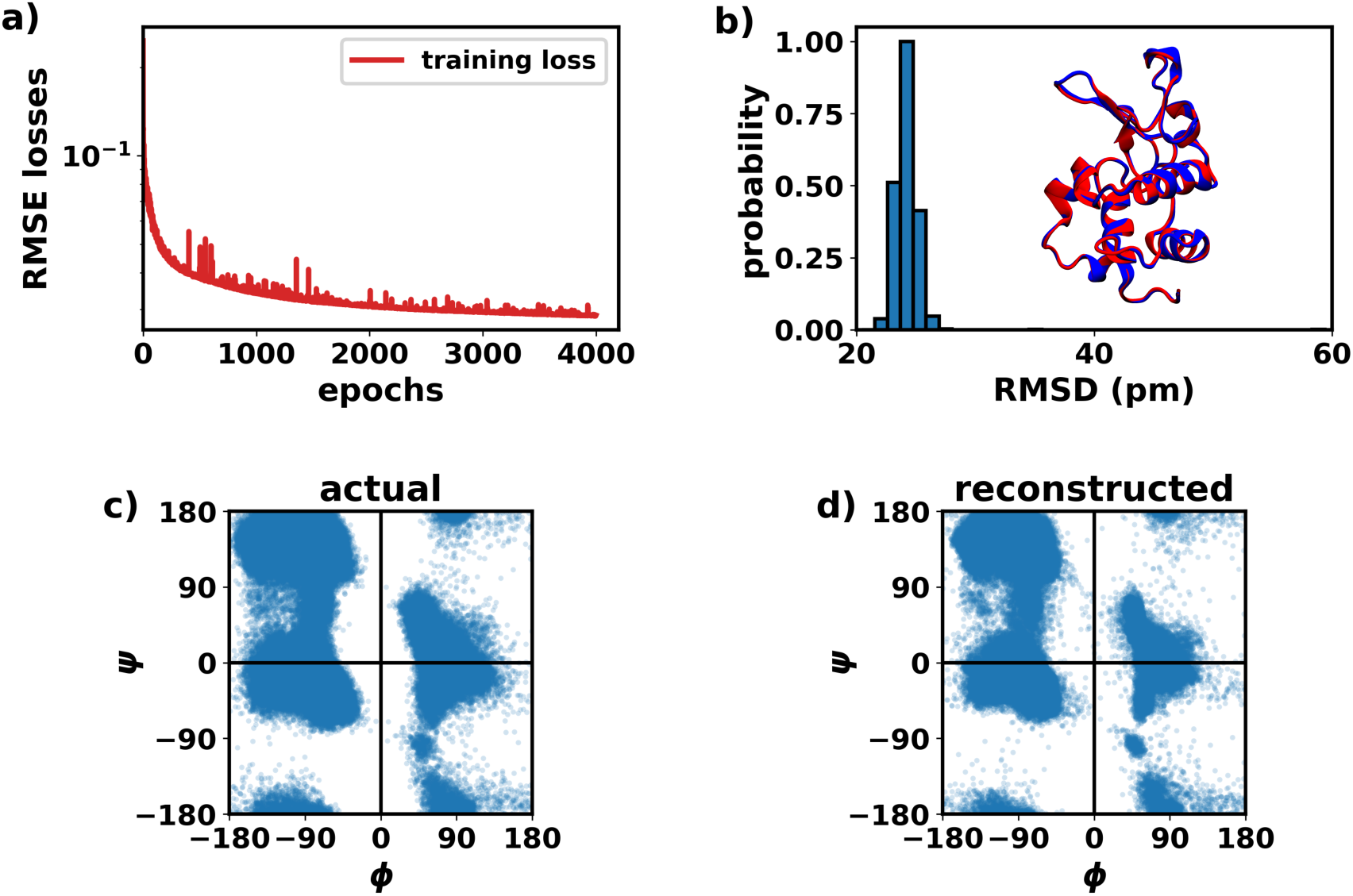
HEWL compression and reconstruction for *c* = 20. **a)** Training loss for HEWL. **b)** Distribution of pairwise root mean squared deviation (RMSD) calculated between original and reconstructed configurations where the original coordinates serve as the reference for the respective reconstructed coordinates. The inset shows the superposition of an original frame and the corresponding reconstructed frame. **c)** Ramachandran plot obtained from original HEWL trajectory. **d)** Ramachandran plot obtained from reconstructed HEWL trajectory.

As the most important result of the entire exercise, we obtain a compression of 85% compared to the corresponding trajectory in the xtc format, which is the – already highly compressed – trajectory file format in GROMACS. The significantly reduced size of the latent space representation of the simulation trajectories, within the range of a few megabytes, makes it very convenient for facile local and cloud storage and for sharing with the community for future exploration of the data.

To verify how reconstruction quality varies with the size of the latent space, we trained AutoEncoders with latent spaces of dimensionality 2, 16, 64, 256, and 1024, while keeping the rest of the AutoEncoder architecture constant. A comparison of the RMSDs for AutoEn-coders trained with different latent spaces is shown in Figure S1a. The results indicate that as the latent space increases, the reconstruction quality improves. However, for large latent spaces (≥ 256), the reconstruction quality reaches an upper limit. The same trend is expected also for other systems, with the latent space dimension beyond which reconstruction quality saturates varying proportionally with system size. Nevertheless, the reconstructions across all latent space dimensions tested are accurate up to the sub-Ångstrøm level.

To check the robustness of the protocol concerning the selected reference frame used to align the trajectories, we trained four models for HEWL using different, randomly chosen reference conformations. Figure S1b shows that the training of models with the same architecture for the same system is independent of the choice of reference conformation.

### Assessment of the automation of the protocol on a diverse range of proteins

To test whether the protocol can be generally applied and automated, we expanded the scope to a set of diverse proteins, ranging from chignolin, trpcage, A*β*40, and ACTR. We fixed the number of latent dimensions to 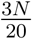 (abbreviated as *c* = 20) and standardized the neural network architecture. For each case, the corresponding network was trained for 1000 epochs. Figures 4a and 4b display the training and validation losses, demonstrating that the networks train effectively using the aligned, centered, and scaled coordinates.

**Figure 4:**
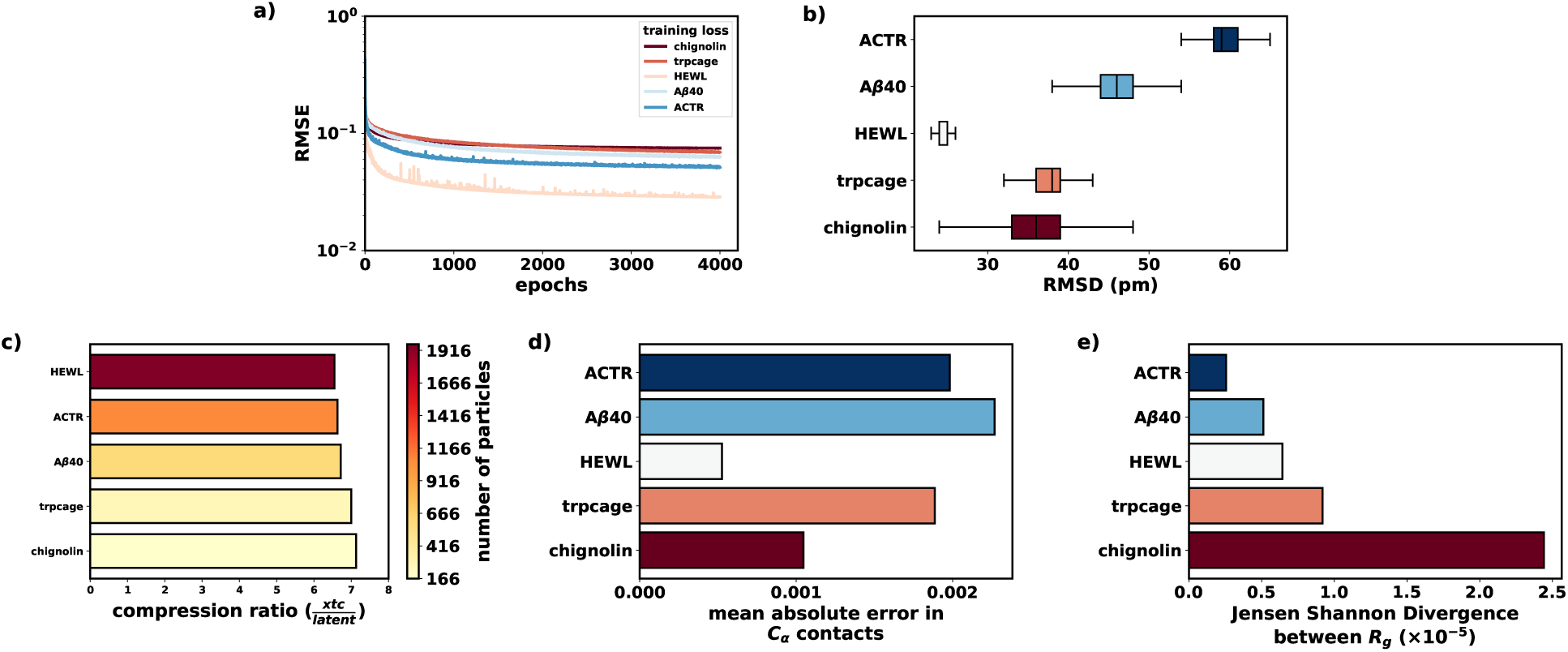
Results for for folded and intrinsically disordered proteins. **a)** Training losses. **b)** Boxplots depicting pair-wise root mean squared deviations (RMSDs). **c)** Extent of compression for *c* = 20. **d)** MAE of differences in residue-residue *C_α_* distances between original and decompressed trajectories. **e)** Jensen Shannon Divergence (JSD) values between *R_g_* distributions obtained from the original and the reconstructed trajectories.

The reconstruction is slightly more accurate for systems with folded proteins compared to intrinsically disordered proteins (IDPs). This difference likely arises because IDPs have pronounced conformational fluctuations, which can affect the alignment of the trajectory with the reference conformation and thus pseudo-SE(3) invariance. Nonetheless, we found that the protocol can still be effectively used for IDPs but with a slightly higher error in reconstruction. The mean RMSD between original and reconstructed trajectory ranges between 10 and 60 pm.

Figure 4b shows boxplots of the distributions of the RMSD values between pairs of corresponding frames from the original and reconstructed trajectories. The low RMSD values indicate that the reconstructed coordinates closely match the original ones.

Figure 4c illustrates the extent of compression for different systems, with the color bar representing the number of particles in each system. The same extent of compression (7 fold or 85%) is reached because *c* = 20 was used for all systems.

To investigate the reconstruction of secondary structures, we used mean absolute errors (MAEs) between original and reconstructed residue-residue C*_α_* contact maps (Figure 4d). As expected from the sub-Ångstrøm precision of the reconstructed atomic coordinates (see above), the MAE values are very low (∼ 10*^−^*^3^), confirming that the backbone structure is accurately reconstructed upon decompression. This is also supported by comparisons of Ramachandran plots between original and reconstructed trajectories (Figure S2). Furthermore, the Jensen-Shannon divergence (JSD) between radius of gyration (*R_g_*) distributions of the original and reconstructed trajectories was used as a measure of the quality of reconstruction of the spatial extension of the protein (Figure 4e). A JSD value of 0 means that two distributions are identical, while a value of 1 corresponds to two disjoint probability distributions. Figure 4e shows that most JSD values are in the order of 10*^−^*^5^, demonstrating accurate reconstruction of the spatial extension(s) of the protein.

### Extension of the protocol to membranes

The above results demonstrate that AutoEncoders can effectively compress and decompress trajectories of individual protein molecules. Next, we extended our approach to membrane systems comprised of multiple lipid molecules. To this end, we used a 300 ns trajectory of a phospholipid bilayer composed of 256 DOPC lipids, with coordinates saved to disk every 100 ps (i.e., 3001 frames). The trajectory was obtained from the NMRlipids database^6^ (entry ID 330). Using the same protocol as described above for the protein trajectories, the DOPC bilayer trajectory was compressed by 85% using *c* = 20.

Figure 5a shows the overlay of a decompressed frame (red) and its original counterpart (blue). For the full trajectory, the pairwise (frame versus frame) RMSD ranges between 41–88 pm, indicating a highly accurate reconstruction of the system. To further validate the properties of the decompressed membrane system, we analyzed the *S_CC_* bond order parameters for the original and decompressed trajectories (Figure 5b). The order parameter is reproduced with high accuracy in the decompressed trajectory. We also analyzed the density of the DOPC head groups (Figure 5c) and of the hydrocarbon tails (Figure S3) along the bilayer normal. Both evaluated properties show excellent agreement with the original data, confirming the high-fidelity reconstruction of the system.

**Figure 5:**
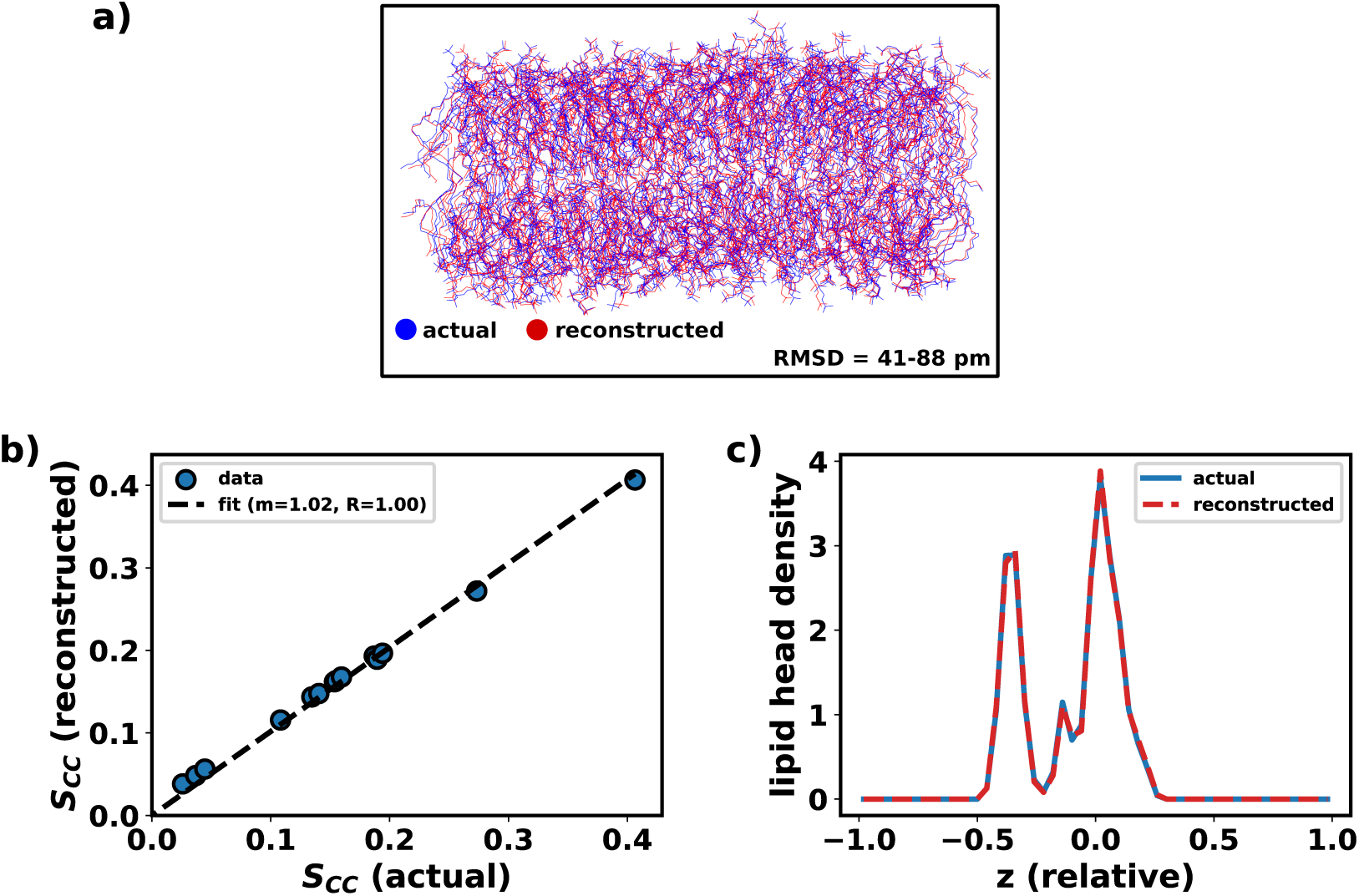
Compression and decompression of a DOPC bilayer. **a)** Superimposed frames from the original (blue) and reconstructed (red) trajectories. The pairwise RMSD range for the full trajectory is between 40–81 pm. **b)** Comparison of *S_CC_* values for the original and reconstructed trajectories. **c)** Densities of the DOPC head groups along the membrane bilayer normal (z-axis of the box).

Taken together, these findings demonstrate that the algorithm is capable of compressing MD trajectories of a phospholipid bilayer and allows for an accurate reconstruction of the original trajectories upon decompression.

### Test on multiple composite systems

We have demonstrated above how AutoEncoders can be effectively used to compress trajectories of single protein systems and lipid bilayers. However, many practical applications involve the simulation of more complex systems, such as protein-ligand or protein-protein complexes, and membrane-bound proteins. Thus, we next demonstrate that the AE algorithm can also be faithfully applied to such complex many-component systems and can reconstruct the relevant functional properties. It should be noted that the ligands’ coordinates would usually occupy only a small amount of disk space, and hence compression of ligand coordinates may not necessarily be required. However, we demonstrate the compression of the ligands as a proof of concept. Additionally, this serves to illustrate how to decouple the coordinates of two (or more) distinct subsystems (here: protein and ligand) before compression and then recouple them after reconstruction.

#### Protein–ligand complexes

For protein–ligand systems, instead of training both protein and ligand in a single neural network, the previously described steps are followed individually for the protein and the ligand. Hence, two AutoEncoders, one for the protein and one for the ligand, are trained and used to produce two compressed latent spaces. To reconstruct the trajectory, the trained decoders are used to decompress the respective protein and ligand trajectories, which are then combined to generate the final reconstructed trajectory (see *Methods*).

We used two systems as test cases, the L99A variant of T4L binding to benzene,^27^ and cytochrome P450 binding to camphor. ^24^ In the first case, the system features a buried hydrophobic pocket in which the symmetric benzene ligand binds via a gate opening event. For cytochrome P450, although it also involves a buried pocket, there is no such gate opening/closing motion. In contrast to benzene, the camphor ligand is asymmetric, and thus it is insightful to check whether the ligand orientations are properly captured after reconstruction. We followed the described protocol to compress and reconstruct the two trajectories. The training losses and the pairwise (frame versus frame) RMSD values indicate accurate reconstruction of the trajectories (Figures 6a and 6b). Using *c* = 20 for protein and *L* = 4 for the ligand, a ∼85% compression is obtained for both systems (Figure 6c). The MAE between residue-residue *C_α_* contacts (Figure 6d) for T4L L99A and cytochrome P450 underlines the accurate reconstruction of the structures.

**Figure 6:**
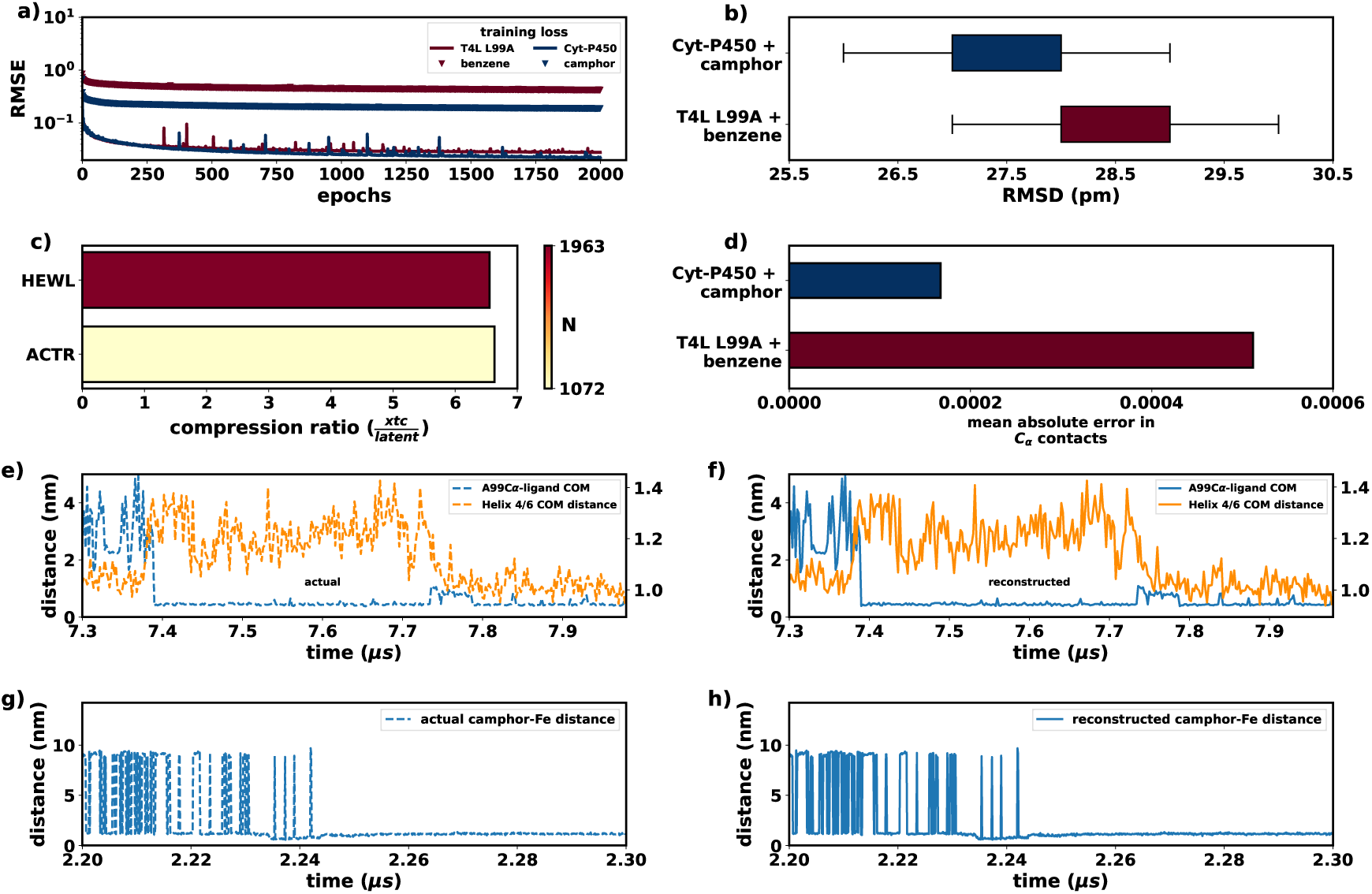
Results for protein-ligand systems. **a)** Training losses for proteins (solid lines) and ligands (triangle markers). **b)** Pairwise RMSD distribution for full systems (protein + ligand). **c)** Compression ratio. **d)** MAE of residue-residue *C_α_* contacts. **e)** The binding profile of benzene to T4L L99A from the original trajectory. **f)** The corresponding ligand binding profile from the reconstructed trajectory. **g)** Binding profile of camphor to cytochrome P450 from the original trajectory. **h)** The corresponding binding profile from the reconstructed trajectory.

##### (i) T4L L99A and benzene

To visualize the accurate reconstruction from the compressed trajectory, Figure S4 shows a superposition of the same frame from the original and the reconstructed trajectories. The overall protein conformation and the ligand positions are properly maintained. We thus next compared the protein dynamics and the ligand binding events between the original and reconstructed trajectories (Figure 6e-f). The reconstructed trajectory accurately describes the original ligand binding event, demonstrating that the extended compression protocol is effective and accurate also for protein-ligand systems.

##### (ii) Cytochrome P450 and camphor

Figure S5a shows a superposition of two corresponding frames from the original and the reconstructed trajectories. As for the T4L/benzene system, the structure of the protein and the position and structure of the ligand are properly maintained. Reconstruction of the binding process of camphor to cytochrome P450 from the compressed trajectory (Figures 6g and 6h) shows accurate reconstruction of the system dynamics. The camphor RMSD distribution and the superimposed ligand conformations (Figure S5b) show that the coordinates are almost exactly reproduced, indicating that the orientation and stereochemistry of the ligand in the protein pocket are preserved.

#### SARS-CoV-2 spike glycoprotein

Next, to also investigate a large composite system with glycosylated amino acids, we turned to the SARS-CoV-2 spike glycoprotein head. The trajectories used in this study were obtained from the Covid-19 BioExcel database^2^ and have been recently published.^25^ Replica 1 was selected for analysis. The system consists of a total of 63,156 atoms (including hydrogens). The full trajectory comprises 50,000 frames, representing a total of 1 *µ*s of simulation time. For testing purposes, we extracted every tenth frame from the trajectory (the original xtc file size is 11 GB).

Application of the compression algorithm (as described in *Methods*) reduced the file size by 85% (relative to the original xtc file). The reconstructed frames exhibit pairwise RMSD ranges of 53 - 110 pm and 82 - 162 pm for the protein and the glycans, respectively. Figure 7a illustrates a reconstructed configuration (red) superimposed with the corresponding frame from the original trajectory (blue); Figure 7b shows a zoom-in on the glycans. These results demonstrate that the algorithm achieves high levels of compression and precise reconstructions after decompression also for very large and complex composite systems such as (glycosylated) protein-protein complexes.

**Figure 7:**
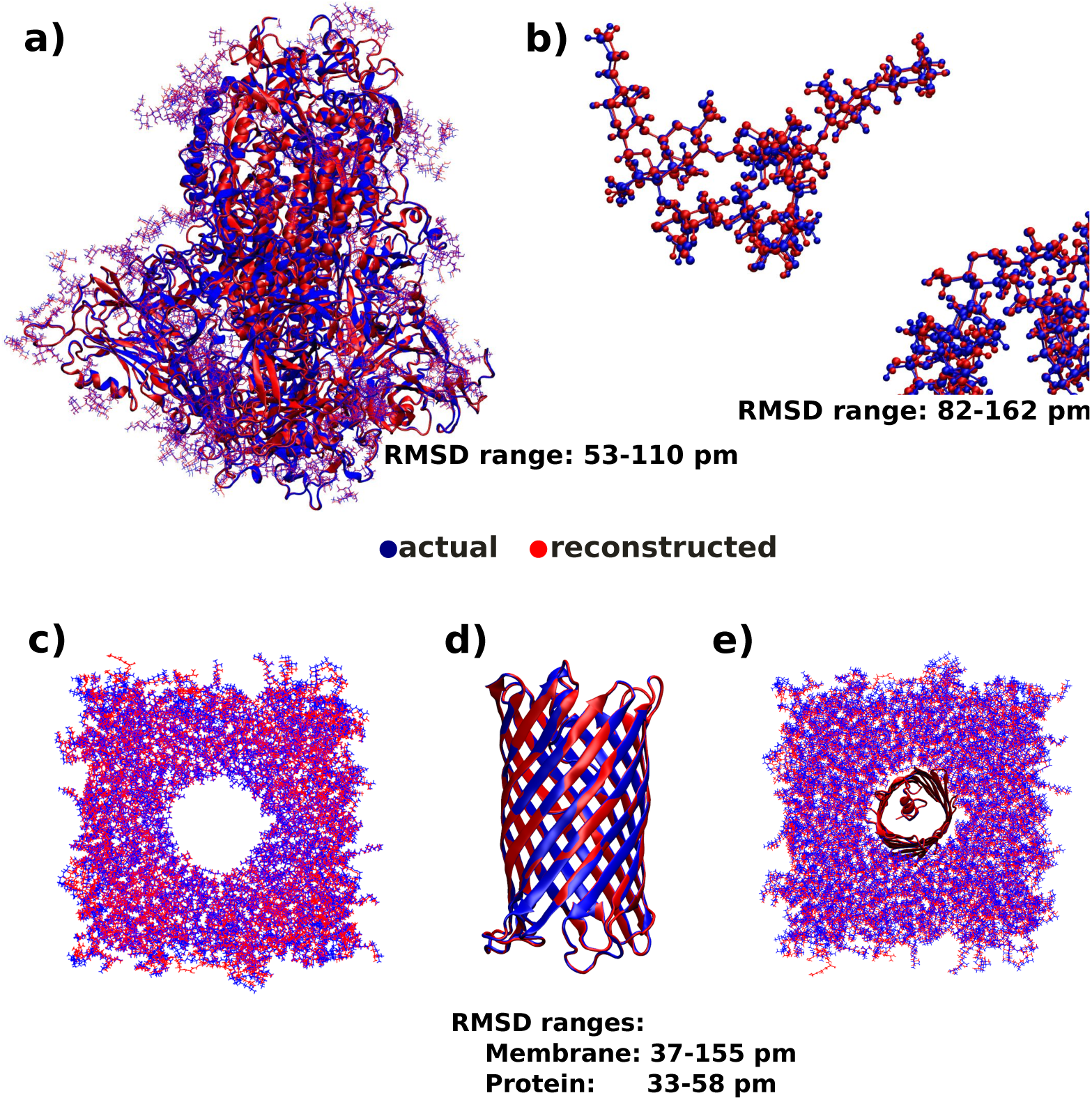
Comparison of original and reconstructed frames of SARS-CoV2 spike protein and membrane-bound CymA. The same frame from the original trajectory and the reconstructed trajectory are shown in blue and red respectively. **a)** The protein with glycans. **b)** Zoom-in on the glycans. The overall pairwise RMSD range is 20-33 pm. **c)** PHPC membrane **d)** CymA transmembrane protein. **e)** PHPC bilayer with embedded CymA. The overall pairwise RMSDs for the membrane and CymA are 40-113 pm and 14-32 pm, respectively.

#### Membrane-bound CymA

We next tested the compression of a membrane-bound bacterial transporter, CymA. The trajectory was obtained from.^26^ The system comprises of the CymA transporter embedded in a PHPC bilayer with lipids in each monolayer, resulting in a total of 40,625 atoms (including hydrogens). To compress this system, we followed the protocol described above for proteinligand systems. Decoupling the protein and the lipids yielded two separate trajectories, one for the protein and one for the membrane. Each trajectory was compressed individually using the protocols described above, resulting in a reduction of approximately 85% in file size.

After compression, the protein and membrane were separately decompressed and then recoupled to reconstruct the full system. The decompressed system exhibits pairwise (frame versus frame) RMSD ranges of 33-58 pm and 37-155 pm for the protein and membrane, respectively (Figures 7c and 7e). We thus conclude that the algorithm can be applied to membrane-bound protein systems as well.

### Compressing liquid water

Finally, to challenge the proposed AE-based compression/reconstruction algorithm with a highly dynamic and diffusive system, we applied it to liquid water. MD simulations in the isobaric-isothermal ensemble (constant temperature of 303.15 K and pressure of 1 bar) were performed for 141 ns with a 4 nm cubic box of TIP3P water, saving coordinates every 20 ps. The resulting trajectory contained 7050 frames.

The standard protocol as described above for single proteins was applied to this system, with the latent space determined by *c* = 20, yielding a reduction of approximately 85% in file size. The pairwise RMSD ranged between 2.3 - 3.1 Å. Figure S6a shows a snapshot of the original trajectory superimposed with the reconstructed frame. It is apparent from Figure S6a that the model does not perform well in reconstructing water. For a quantitative understanding of the model’s performance, we calculated the oxygen-oxygen radial distribution functions (RDF) for the actual and the reconstructed trajectories, which are compared in Figure S6b. Figure S6b shows major discrepancies between the actual and the reconstructed oxygen-oxygen RDFs. Therefore, we conclude that one should be cautious with applying the current compression/reconstruction protocol to liquid water, or probably to liquids in general.

### Reconstruction quality versus compression

We demonstrated above that the AE-based latent space compression method can yield a significant reduction of file sizes of MD simulation trajectories, with consecutive decompression providing high-accuracy reconstructions of the original atomic coordinates. Although one can achieve compression levels of up to 85% with reconstruction RMSDs as low as a few picometers, users may still be concerned about potential amino acid stereochemistry issues that might not be adequately captured by the RMSD. More importantly, the protocol should be able to provide quasi-exact reconstruction if needed, even if this requires sacrificing some degree of compression.

To demonstrate the robustness of the protocol, we trained 5 AE models on T4L L99A (protein only), each having a different latent space dimension. For each model, we compressed and subsequently decompressed the trajectory. From the resulting trajectories, we randomly selected 5 representative pairs of frames, where each pair consisted of a frame from the original trajectory and the corresponding frame after compression and decompression. This resulted in 25 frame pairs, for which we calculated MolProbity scores.^46^ Figure S7 shows the variation of MolProbity scores with latent space size for the reconstructed frames. For *L* = 1024 (60% compression), the score drops below 2.2, indicating that the reconstructed structures are of very high quality. We also observed that the chiral centers of the amino acids, along with other stereochemical features, are fully preserved. A detailed comparison of multiple such metrics, including those discussed above, is summarized in *Supplementary File 1*, where the suffix 0 indicates values from the original frames.

Finally, going beyond atomic positions, we calculated the potential energies of the original and reconstructed configurations with the CHARMM36m force field (*Supplementary File 2*). We observed that, in a few cases, the potential energy of the reconstructed configuration was high, primarily due to the Lennard-Jones contribution, which indicates atomic clashes. As a proof of concept, we selected the reconstructed frame with the highest energy (L = 2048, replica = 2), solvated it in TIP3P water, and added ions for charge neutralization using CHARMM-GUI.^47^ Energy minimization of the system to a maximum force tolerance threshold of 1000 kJ mol*^−^*^1^ nm*^−^*^1^ was achieved in 803 steepest descent steps. The super-imposed conformations (Figure S8) show that the energy-minimized conformation and the reconstructed conformation are similar, with an RMSD of 0.3 Å. The high potential energy is mainly due to the steeply repulsive (*r*^−12^) nature of the Lennard-Jones potential at short interatomic distances and also because of the stiff bond and angle potentials, so that even very small positional deviations can lead to large increases in energy. Thus, energy minimization should be performed before using the reconstructed configurations as starting points for MD simulations.

## Conclusions

In this study, we demonstrated the use of AutoEncoders (AEs) to very effectively compress Molecular Dynamics (MD) simulation trajectories. By developing a protocol that involves choosing a reference frame, aligning the trajectory, and scaling the coordinates, precise reconstruction of atomic coordinates was achieved. Our approach yielded high accuracy in reconstructing the original data across various test cases by employing a consistent AutoEn-coder architecture, trained for at least 2000 epochs (depending on the system).

We validated our protocol by comparing multiple structural and functional properties calculated from the original and decompressed trajectories. The reconstructed data closely match the original trajectories. This has been demonstrated by the reconstructed trajectories, which successfully recapitulate a range of structurally and functionally relevant biophysical properties. In particular, the protocol accurately reproduces key features of the protein’s Ramachandran map, C*_α_* contacts, radii of gyration, as well as lipid order parameters and density profiles. We also demonstrated that the protocol can be seamlessly used for protein-ligand complexes, large protein-protein complexes and mixed membrane/protein systems.

Our AutoEncoder (AE)-based method advances the state of the art by learning a compact, non-linear representation of trajectory frames, eliminating the need for handcrafted predictors or domain-specific tuning. The protocol is system-agnostic and results in ≥85% file size reduction over the already highly compressed xtc format, while maintaining sub-Å positional accuracy. By training a neural network to extract latent representations across diverse motions, our method captures both local and global correlations, including complex dynamics that are challenging for linear or rule-based methods. Despite achieving high compression, it reconstructs biomolecular trajectories with high fidelity, preserving essential structural features with RMSD values in the sub-Å range for proteins and lipids, across timescales ranging from hundreds of nanoseconds to several microseconds.

Our protocol enables efficient storage and sharing of MD simulation data, facilitating broader exploration and utilization of existing datasets. The robustness, flexibility, and high accuracy of our method make it a valuable tool for the scientific community, especially for storing and distributing large-scale biomolecular simulation data. We anticipate that such AE-based approaches can also be employed for compressing other large datasets, e.g., from cryoEM, which currently pose significant challenges due to their size.

## Code and Data Availability

The codes, along with examples, can be found at https://github.com/SerpentByte/compresstraj. Most of the MD data of the test systems used have been obtained from D.E. Shaw Research with permission.^20^ The data for T4L L99A + benzene were obtained from Ref.^27^ The simulation trajectories for cytochrome P450 + camphor were obtained from Ref.^23^ and a smaller version was used as an example. The trajectory for SARS-CoV-2 spike protein was obtained from Refs.^2,25^ and for membrane-bound CymA from Ref.^26^

## Supporting Information

The Supporting Information (SI) provides supplemental figures, supplemental table. The SI Methods contains details on the architecture of the AutoEncoder used. Figure S1 shows the variation of RMSD with the latent space and also with the reference configuration used. Figure S2 compares the Ramachandran plots between the actual and the reconstructed trajectories. Figure S3 compares the lipid tail densities along the bilayer normal between actual and reconstructed trajectories. Figure S4 compares the reconstruction of the protein for T4L L99A + benzene. Figure S5 compares the reconstruction of Cytochrome P450 + camphor. Figure S6 compares the reconstruction of a 300 ns long water trajectory. Figure S7 shows that with increases latent space, MolProbity scores monotonically decreases untill it reaches a minima. Figure S8 compares a reconstructed conformation with the highest potential energy with its energy minimized version. SI File 1 contains the data required to plot Figure S7. SI File 2 contains the comparison of energies between actual frames and their reconstructed counterparts.

## Acknowledgement

This study was carried out during the visit of AW and JM in the group of LVS at Ruhr University Bochum. We acknowledge the funding by the Alexander von Humboldt-Stiftung through an Experienced Researcher Fellowship that supported J.M.’s visit. We acknowledge support of the Department of Atomic Energy, Government of India, under Project Identification No. RTI 4007. This work was supported by the Deutsche Forschungsgemeinschaft (DFG) under Germany’s Excellence Strategy - EXC 2033 - 390677874 - RESOLV.

