## supplementary_information for "Employing Artificial Neural Networks for Optimal Storage and Facile Sharing of Molecular Dynamics Simulation Trajectories"

### Supplementary Methods

#### Details on the AutoEncoder neural network layers

In practice N is substituted by the number of particles present in the system multiplied by 3 and L is substituted by the desired latent space dimensions.

```
DenseAutoEncoder
|---- Encoder
|    |---- Linear(in_features=N, out_features=4096, bias=True)
|    |---- BatchNorm1d(4096, eps=1e-05, momentum=0.1, affine=True, track_running_stats=True)
|    |---- ELU(alpha=1.0)
|    |---- Linear(in_features=4096, out_features=1024, bias=True)
|    |---- BatchNorm1d(1024, eps=1e-05, momentum=0.1, affine=True, track_running_stats=True)
|    |---- ELU(alpha=1.0)
|    |---- Linear(in_features=1024, out_features=L, bias=True)
|    |---- ELU(alpha=1.0)
|---- Decoder
|    |---- Linear(in_features=L, out_features=1024, bias=True)
|    |---- BatchNorm1d(1024, eps=1e-05, momentum=0.1, affine=True, track_running_stats=True)
|    |---- ELU(alpha=1.0)
|    |---- Linear(in_features=1024, out_features=4096, bias=True)
|    |---- BatchNorm1d(4096, eps=1e-05, momentum=0.1, affine=True, track_running_stats=True)
|    |---- ELU(alpha=1.0)
|    |---- Linear(in_features=4096, out_features=N, bias=True)
|    |---- Sigmoid()
```

### Supplementary Figures

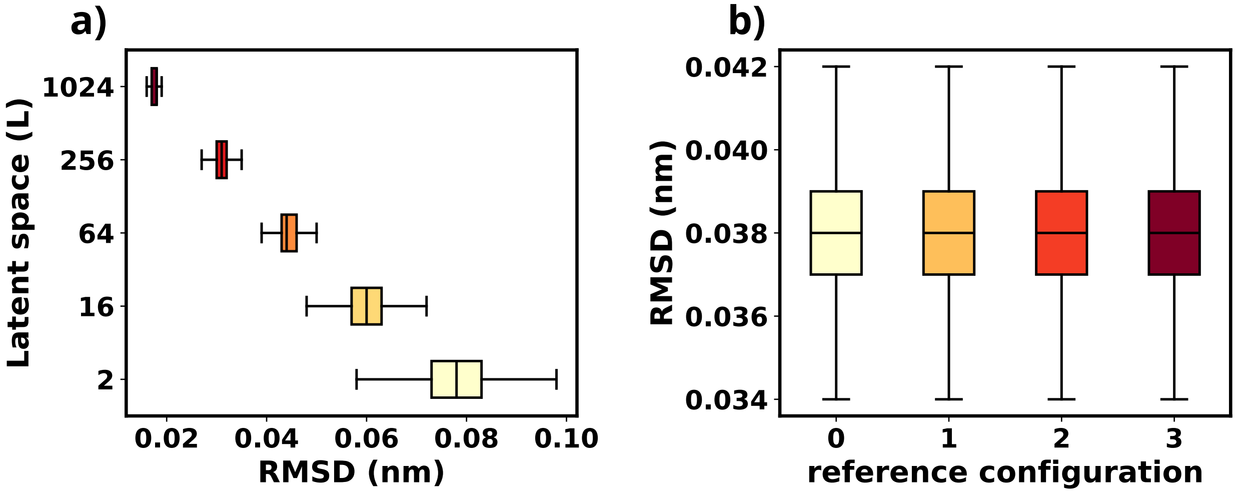


Figure S1: **a)** Variation of pairwise RMSD of reconstructed structures for different latent space dimensions. *b)* RMSD variation during trajectory reconstruction using different reference frames for alignment.

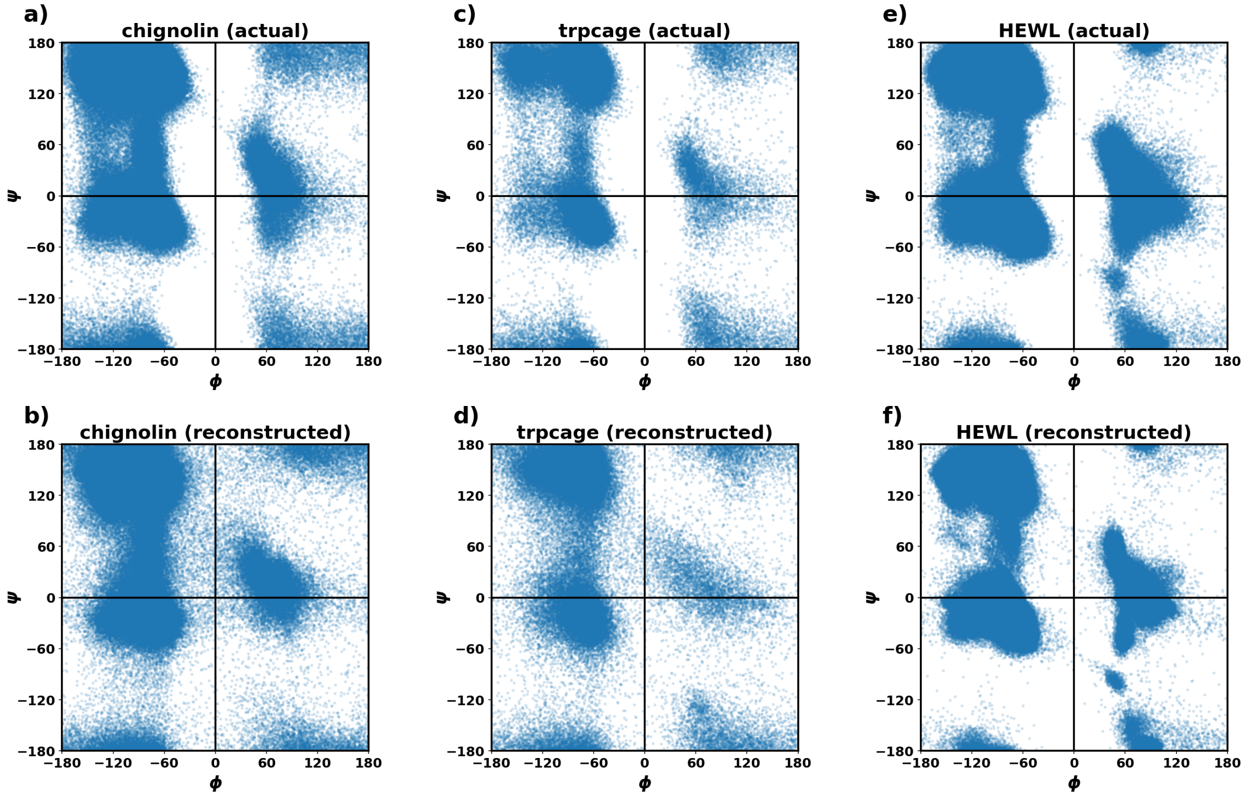


Figure S2: Comparison of Ramachandran plots obtained from original and reconstructed trajectories for different folded proteins.

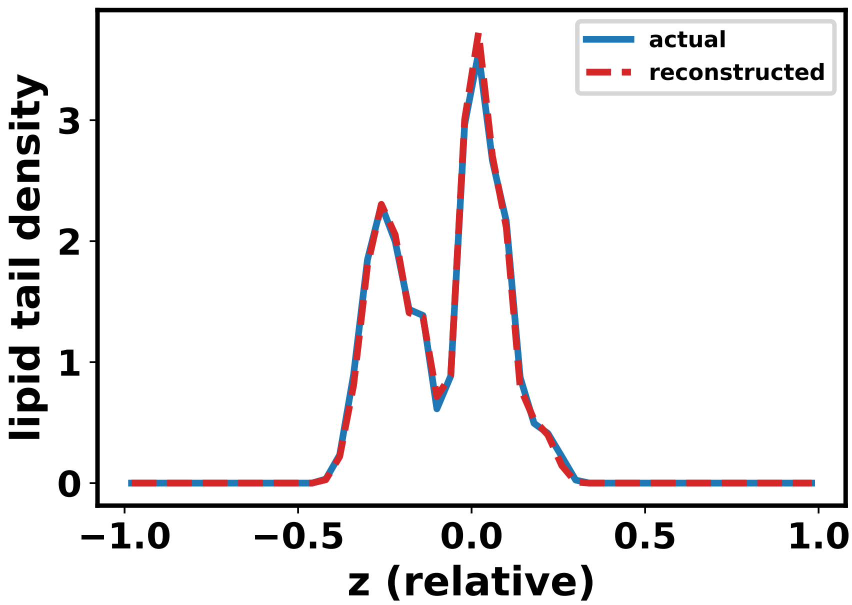


Figure S3: Density of DOPC tail carbons along the bilayer normal (z-axis of the box).

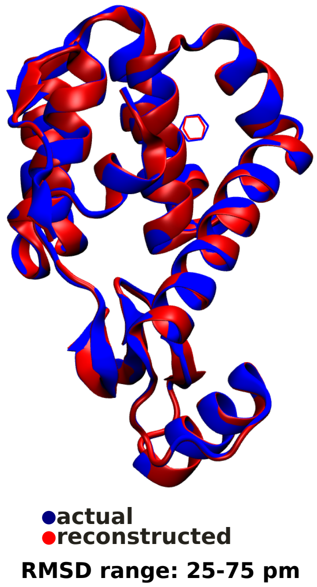


Figure S4: **a)** Superposition of the same frame from the original (blue) and the reconstructed (red) trajectories of T4L L99A + benzene.

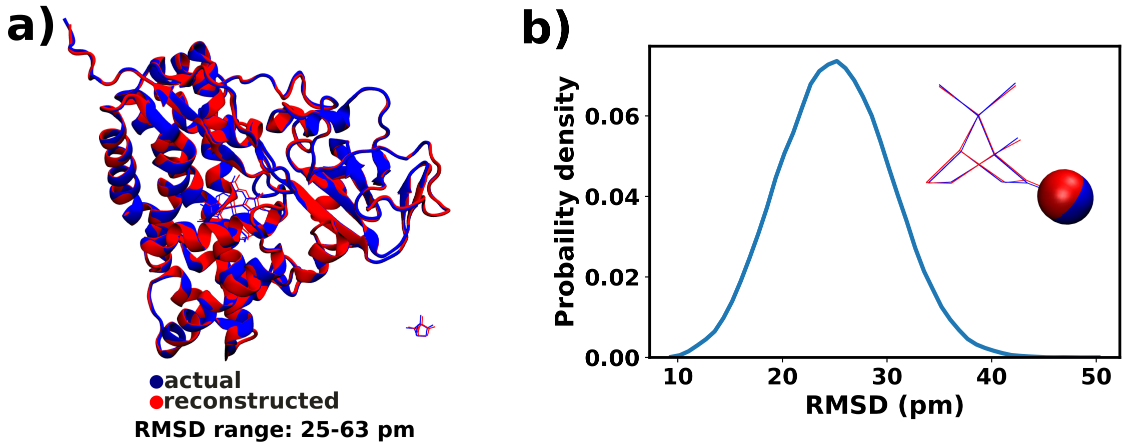


Figure S5: **a)** Superposition of the same frame from the original (blue) and the reconstructed (red) trajectories of Cytochrome P450 + camphor. **b)** Pairwise RMSD distribution for camphor. The inset shows a superposition of camphor conformations from the same frame but obtained from the original (blue) and the reconstructed (red) trajectories. Oxygen is shown as a red sphere and carbons and the backbone are colored black. Hydrogens are not shown.

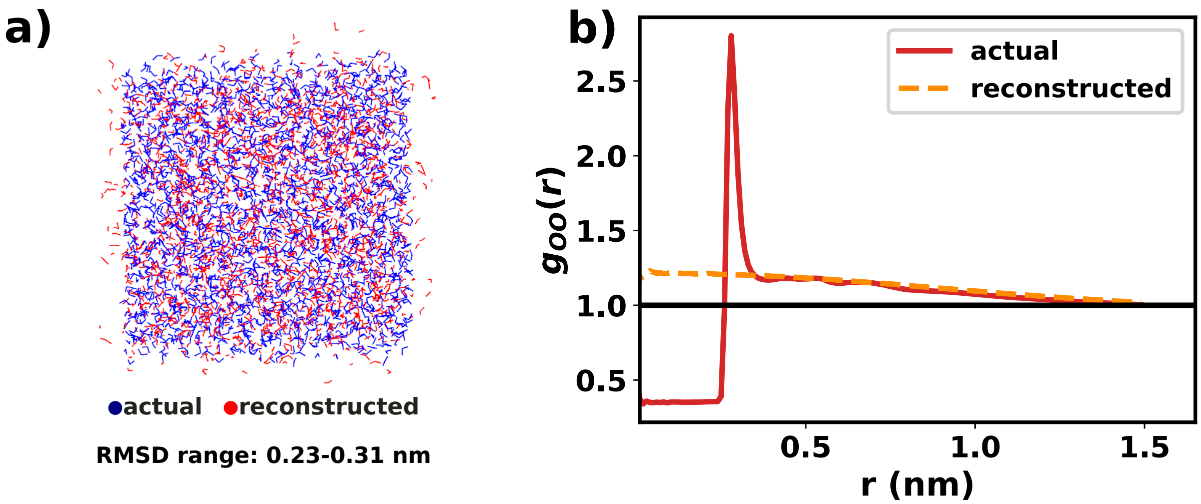


Figure S6: **a)** Comparison of original and reconstructed trajectories of TIP3P water. The same frame from the original trajectory and the reconstructed trajectory is shown in blue and red respectively. **b)** Comparison of oxygen-oxygen radial distribution functions from original and reconstructed water trajevtories.

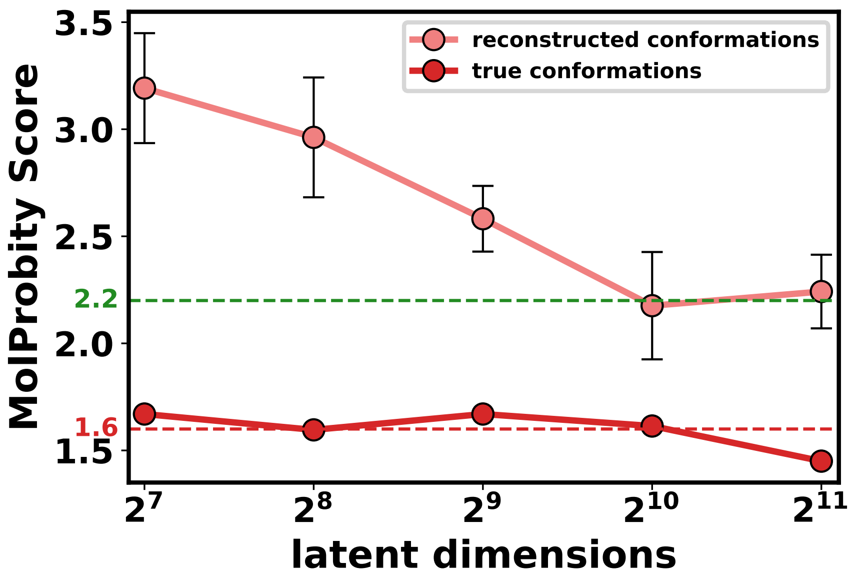


Figure S7: MolProbity score[1] variation with the latent space dimensions for T4L L99A. The green dashed line shows a threshold below which structures are typically considered to have super high fidelity. The red dashed line indicates the average scores of the original structures. The light red points/line represent the MolProbity scores of the reconstructed structures.

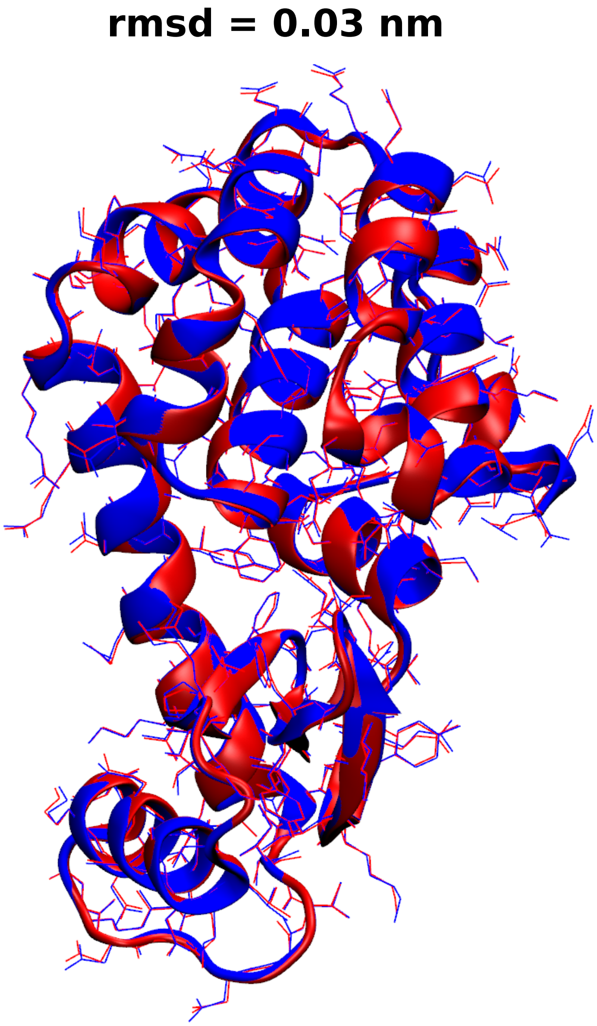


Figure S8: Superimposition of the following conformations for T4L L99A: **red** represents the reconstructed structure with the highest potential energy (L=2048, replica=2); **blue** represents the same conformation after energy minimization using CHARMM36m, in the presence of water and ions required for charge neutralization. The two conformations have an RMSD of 0.3 Å.
